# QUANTIFYING INTRASPECIFIC VARIATION IN HOST RESISTANCE AND TOLERANCE TO A LETHAL PATHOGEN

**DOI:** 10.1101/2023.09.29.557180

**Authors:** Bennett M. Hardy, Erin Muths, W. Chris Funk, Larissa L. Bailey

## Abstract

Due to the ubiquity of disease in natural systems, hosts have evolved strategies of disease resistance and tolerance to defend themselves from further harm once infected. Resistance strategies directly limit pathogen growth, typically leading to lower infection burdens in the host. A tolerance approach limits the fitness consequences caused by the pathogen but does not directly inhibit pathogen growth. Testing for intraspecific variation in wild host populations is important for informing conservation decisions about captive breeding, translocation, and disease treatment. Here, we test for the relative importance of tolerance and resistance in multiple populations of boreal toads (*Anaxyrus boreas boreas*) against *Batrachochytrium dendrobatidis* (Bd), the amphibian fungal pathogen responsible for the greatest host biodiversity loss due to disease. Boreal toads have severely declined in Colorado (CO) due to Bd, but toad populations challenged with Bd in western Wyoming (WY) appear to be less affected. We used a common garden infection experiment to expose post-metamorphic toads sourced from four populations (2 in CO and 2 in WY) to Bd and monitored changes in mass, pathogen burden, and survival for eight weeks. We used a multi-state modeling approach to estimate weekly survival and transition probabilities between infected and cleared states, reflecting a dynamic infection process that traditional approaches fail to capture. We found that WY boreal toads are highly tolerant to Bd infection with higher survival probabilities than those in CO when infected with identical pathogen burdens. WY toads also had higher probabilities of clearing infections and took an average of five days longer to reach peak infection burdens. Our results demonstrate strong intraspecific differences in tolerance and resistance that explain why population declines vary regionally across the species. We used a robust, multi-state framework to gain inference on typically hidden disease processes when testing for host tolerance or resistance and demonstrated that describing an entire species as ‘tolerant’ or ‘resistant’ is unwise without testing for intraspecific variation in host defenses.

## INTRODUCTION

Disease strongly affects all scales of biological organization, from influencing individual behavior (e.g., Buck et al. 2018) and population dynamics (Hudson et al. 1998), to structuring communities (Wood et al. 2007), affecting whole ecosystems (Monk et al. 2022), and ultimately shaping the evolution of both host and pathogen (May and Anderson 1983). Due to the ubiquity of disease in natural systems, hosts have evolved strategies to defend themselves from infection, or further harm once infected. These defenses fall into two general categories: 1) mechanisms of disease resistance, and 2) mechanisms of disease tolerance. Host resistance mechanisms are defined as those that directly affect the pathogen and limit its growth (Råberg et al. 2009). For example, skin or gut microbiota that naturally occur in many organisms are responsible for limiting infections by controlling pathogen growth and are considered a mechanism of disease resistance (Van Den Elsen et al. 2017). Host tolerance mechanisms are defined as those that limit the damage or negative fitness consequences caused by the pathogen and do not directly inhibit pathogen growth (Råberg et al. 2009). Host cellular and tissue repair responses are examples of tolerance mechanisms that aid in reducing damage caused by infection (Medzhitov et al. 2012). For instance, mice with a specific enzyme were able to tolerate bacterial infections because the enzyme helped prevent tissue damage due to circulating free heme (Larsen et al. 2010). As ecologists are increasingly aware of the growing contribution of emerging infectious diseases to the loss of biodiversity (Daszak et al. 2000; Smith et al. 2006, 2009), investigating whether host resistance or tolerance exists in wildlife species of concern could be particularly important for predicting host-pathogen dynamics and informing conservation actions.

Distinguishing between resistance or tolerance mechanisms in host populations can inform our expectations for evolutionary dynamics of both host and pathogen and guide conservation strategies. Because host resistance mechanisms impact pathogens negatively by limiting their growth, host resistance may select for pathogens that can overcome resistance mechanisms due to shorter pathogen generation times than hosts, thus quickly evolving and leading to antagonistic coevolution (Roy and Kirchner 2000). The presence of resistance traits in a host population is predicted to fluctuate via a negative feedback loop shaped by the proportion of resistant individuals in a population and how beneficial the trait is (Roy and Kirchner 2000). Because maintaining resistance is assumed to have an energetic cost, selection will act against hosts when pathogen prevalence is nearly absent, such that the cost of maintaining resistance traits is relatively higher when their advantage is minimal (Roy and Kirchner 2000). Thus, resistance strategies may be optimal for hosts where infection risk is transient or unpredictable (e.g., pathogen seasonality or vector driven) and in large, diverse host communities that exist along a continuum of susceptibility, where several resistant hosts will benefit those that are less resistant. If resistance is identified in a given host-pathogen system, this may signal that conservation actions aimed at assisting pathogen clearance or eradication will be beneficial (e.g., increasing environmental features that aid in host clearance).

In contrast, tolerance mechanisms do not impact the pathogen negatively and pathogen prevalence is expected to increase as tolerant hosts increase. While increased tolerance may seem like a superior host strategy compared to resistance, the increased emergence of tolerant reservoir hosts may amplify and promote pathogen persistence and could aid in increased transmission to more vulnerable, intolerant hosts. Tolerance mechanisms therefore may be optimal host strategies where risk of infection is relatively constant through space or time (e.g., environmental persistence or environmental transmission of the pathogen), and where susceptible host communities are less diverse or exist at lower densities to minimize the effect of potential tolerant superspreaders to others within the community (Råberg et al. 2009). If host tolerance is identified, this might signal that pathogen prevalence or infection risk is relatively high, and that measures aimed at eradicating the pathogen may be ineffective. Though resistance and tolerance are presented as a dichotomy, it is important to note that these host defense strategies are not mutually exclusive, and both can coexist within a single host population (Pagán and García-Arenal 2018). The differences in host-pathogen dynamics described above ultimately illustrate the importance of identifying variation in host resistance and tolerance in natural systems as each strategy begets a different outcome for host and pathogen.

Host tolerance or resistance is a major component in determining whether hosts persist or perish in the face of disease (Brannelly et al. 2021; Russell et al. 2020). Regardless of which defense strategy is present, accurately describing variation in host tolerance or resistance across the landscape is important for understanding the relative vulnerability of populations to disease. For example, information on population variation for tolerance or resistance could be used to identify potential source populations for supplementation (moving individuals from relatively tolerant or resistant populations to those that are less tolerant or resistant), reintroduction (moving tolerant or resistant individuals to locations that were historically occupied but not currently), and for captive breeding and assurance programs.

Much of the host-susceptibility research using non-model wildlife systems to date have focused on interspecific comparisons of resistance or tolerance and use a single population to represent species-wide patterns (e.g., Woodhams et al. 2007, Sears et al. 2011, Murone et al. 2017, DeSimone et al. 2018, Grab et al. 2019, Knutie et al. 2016, Knutie et al. 2017). These approaches risk over-generalizing a species’ susceptibility to disease, or ability to persist with disease, and could lead to categorizing an entire species as ‘tolerant’ or ‘resistant’, without accounting for the underlying population variation that is likely common. The consequences of ignoring intraspecific variation across biological disciplines has recently gained attention in the study of species traits in evolutionary and community ecology (Bolnick et al. 2011; Des Roches et al. 2018) and impacts of climate change on organismal physiology (Bennett et al. 2019; Cicchino et al. 2023). Continuing the coarse-scale investigation of species’ resistance, tolerance, and susceptibility to pathogens and parasites without assessing intraspecific host variation may also hinder our understanding of host-pathogen dynamics and species conservation.

Researchers have started to investigate intraspecific variation in host tolerance and resistance in wildlife systems using experimental pathogen exposures to hosts, but several limitations restrict the inferences gained from these studies. Often, studies use a single representative population across different strata (e.g., strong and weak demographic responses or high and low disease prevalence) such that there is no replication within strata (e.g., Waddle et al. 2019; Atkinson et al. 2013; Bonneaud et al. 2011; Adelman et al. 2013; Bonneaud et al. 2019). Evidence of host resistance or tolerance using multiple population replicates across strata is uncommon (but see: Savage and Zamudio 2011; Rocke et al. 2012; Henschen et al. 2023; Grogan et al. 2023), but vitally necessary to confidently assess variation in resistance or tolerance. If population replication across strata of interest is not included, results from such studies may lead to missed management opportunities, or even un-mitigated population declines. Furthermore, analyses of resistance or tolerance from experimental exposures are often limited by representing only static, group-level effects on fitness outcomes (e.g., odds of survival from time-to-event analyses; Le-Rademacher et al. 2022). These classical methodologies miss the dynamic processes of disease by not accounting for a host’s transition between various states of interest as disease progresses (e.g., infected, cleared, showing clinical signs). Multistate models provide a flexible approach to estimate parameters of interest in epidemiological studies that include multiple states, events, and measures of dynamic covariates and are commonly employed in various epidemiological field studies of marked hosts (e.g., Cooch et al. 2012; Conn and Cooch 2009; Muths et al. 2020). Their advantages make them particularly attractive to apply to investigations of host tolerance or resistance and are not found in studies of experimental exposures in the context of wildlife disease dynamics.

We use an experimental exposure of boreal toads (*Anaxyrus boreas boreas*) to *Batrachochytrium dendrobatidis* (Bd) across multiple host populations to illustrate the utility of the multistate framework in revealing intraspecific differences in host tolerance and resistance to pathogen infection. Bd is a fungal pathogen contributing to global amphibian declines and extirpations (Skerratt et al. 2007; Wake and Vredenburg 2008) and is considered responsible for the largest loss of host biodiversity due to a single pathogen (Scheele et al. 2019). While many studies have used experimental exposures to describe variation in susceptibility to Bd among amphibian species (Sauer et al. 2020), only a handful have investigated intraspecific variation among populations in susceptibility (Tobler and Schmidt 2010; Savage and Zamudio 2011; Bataille et al. 2015; Waddle et al. 2019). Laboratory studies that examine the relative roles of tolerance vs. resistance using a robust quantitative framework directly linking host fitness and pathogen burden have yet to be implemented broadly (Grogan et al. 2023).

The boreal toad system is ideal for investigating intraspecific variation in host responses to disease and highlights the potential danger in overgeneralizing a species’ response range-wide. Boreal toad populations in the United States have experienced severe declines and extirpations due to Bd in the southern Rocky Mountains region of their range in Colorado and southeastern Wyoming (Hardy et al. 2023; Mosher et al. 2018; Muths et al. 2003). Conversely, populations in western Wyoming and Montana have persisted despite high Bd prevalence within and among populations (Hossack et al. 2020; Pilliod et al. 2010; Muths et al. 2011; Russell et al. 2019). This apparent dichotomy in demographic responses provides an ideal natural laboratory to investigate whether intraspecific variation in resistance or tolerance to Bd exists within boreal toads. We hypothesize that the differences in the relative severity of declines in populations that have been exposed to Bd is due to intraspecific variation in tolerance, resistance, or both. To test this hypothesis, we use a common garden exposure experiment to investigate differences in host fitness when challenged with Bd (Figure 1).

**Figure 1.**
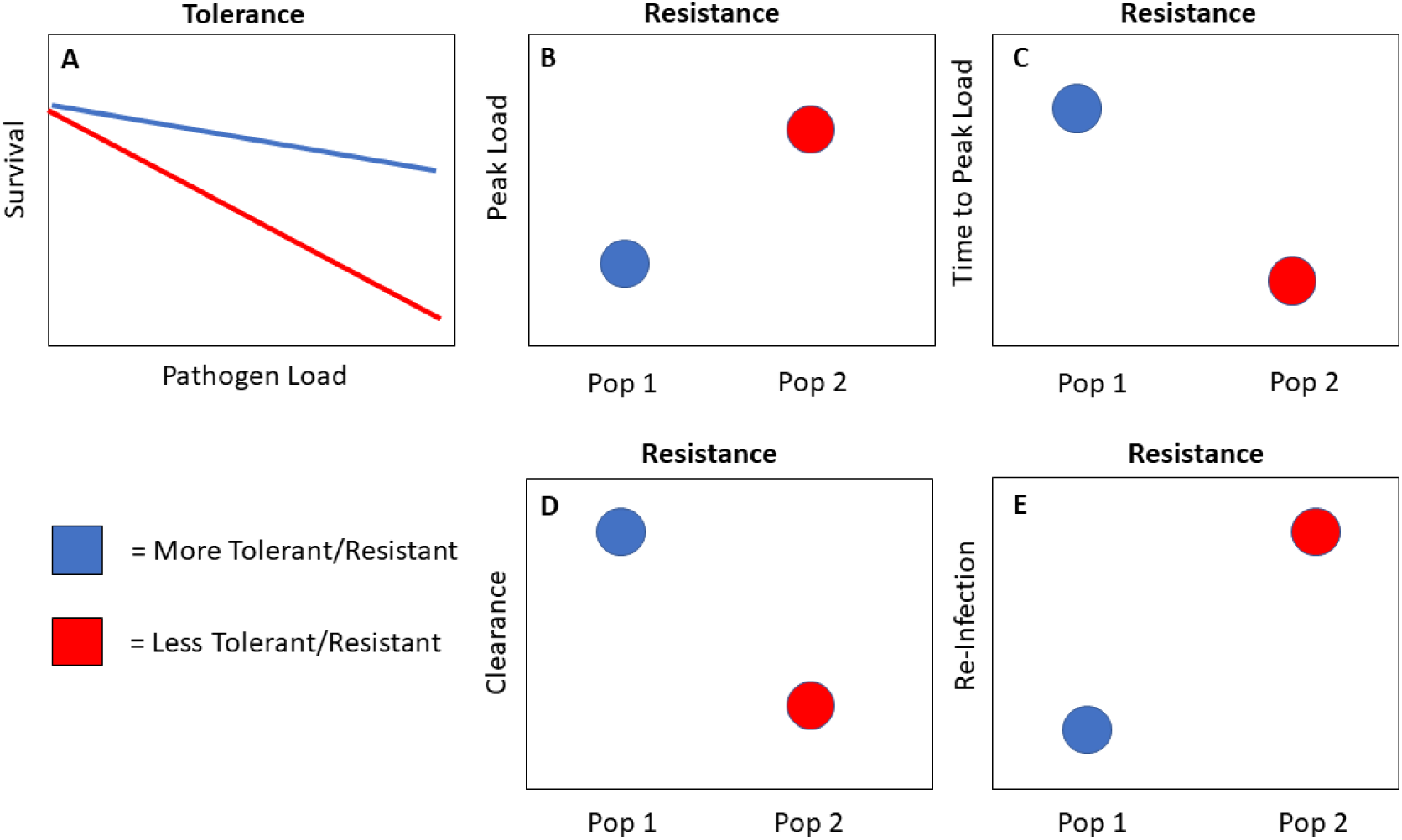
Predictions of intraspecific variation for relative tolerance (Panel A) or resistance (Panels B-E) for two host populations of interest. A. Lines represent the predicted relationship between survival (a metric of fitness) and pathogen load (a measure of burden) for individuals from two populations of differing tolerance. Individuals from the blue population are more tolerant than the red population because, at any given pathogen load, they have higher survival. B. Circles represent population averages for peak pathogen load: population 1 is more resistant than population 2 because, on average, it has the lower peak pathogen load. C. Circles represent population averages for the time (e.g., days, weeks) it takes to reach peak pathogen loads: population 1 is more resistant than population 2 because the pathogen growth is slower, taking longer to reach peak pathogen loads. D. Circles represent mean estimates of the probability of pathogen clearance: population 1 is more resistant because it has higher probabilities of clearing infections. E. Circles represent mean estimates of the probability of re-infection after clearance: population 1 is more resistant because it has lower probabilities of becoming re-infected.

## METHODS

### Experimental design

We selected four boreal toad populations to collect eggs from for use in our exposure experiment (Figure 2). All populations have a history of Bd presence and tested positive for Bd in 2021. We selected two populations on US Forest Service lands in western Wyoming (Chall Creek and Blackrock) and two populations in Colorado (Lost Lake, located in Rocky Mountain National Park, and Panhandle Creek on US Forest Service land). For details on egg, tadpole, and toad husbandry see Appendix 1. When toads from a population averaged 6-7 weeks post-metamorphosis, we transferred them to a temperature-controlled experimental room for individual Bd treatments. During the experiment toads were individually housed in 946ml plastic containers with Eco Earth® (Zoo Med Laboratories Inc., San Luis Obispo, California, USA) substrate, a small dish of water, and a mesh lid for air circulation. Toads were fed flightless *Drosophila melanogaster* every other day and water was re-filled every day. Toads from each collection site were randomly assigned to one of three Bd exposure treatments: low (1×10^2^ zoospores), medium (1×10^3^), or high (1×10^5^). Doses were selected to ensure infections were distributed across a range of zoospore burdens to assess potential differences in tolerance or resistance. Prior to exposure, toads were weighed to the nearest 0.10 gram and swabbed with sterile cotton swabs (Fisher Scientific) following Livo et al. (2004) to verify that toads were Bd-free. We randomly assigned toads to each treatment group and we conducted an ANOVA to confirm that average body mass did not differ among groups. Toad habitats were arranged on metal shelving racks in a randomized block design to avoid potential environmental inconsistencies within the room.

**Figure 2.**
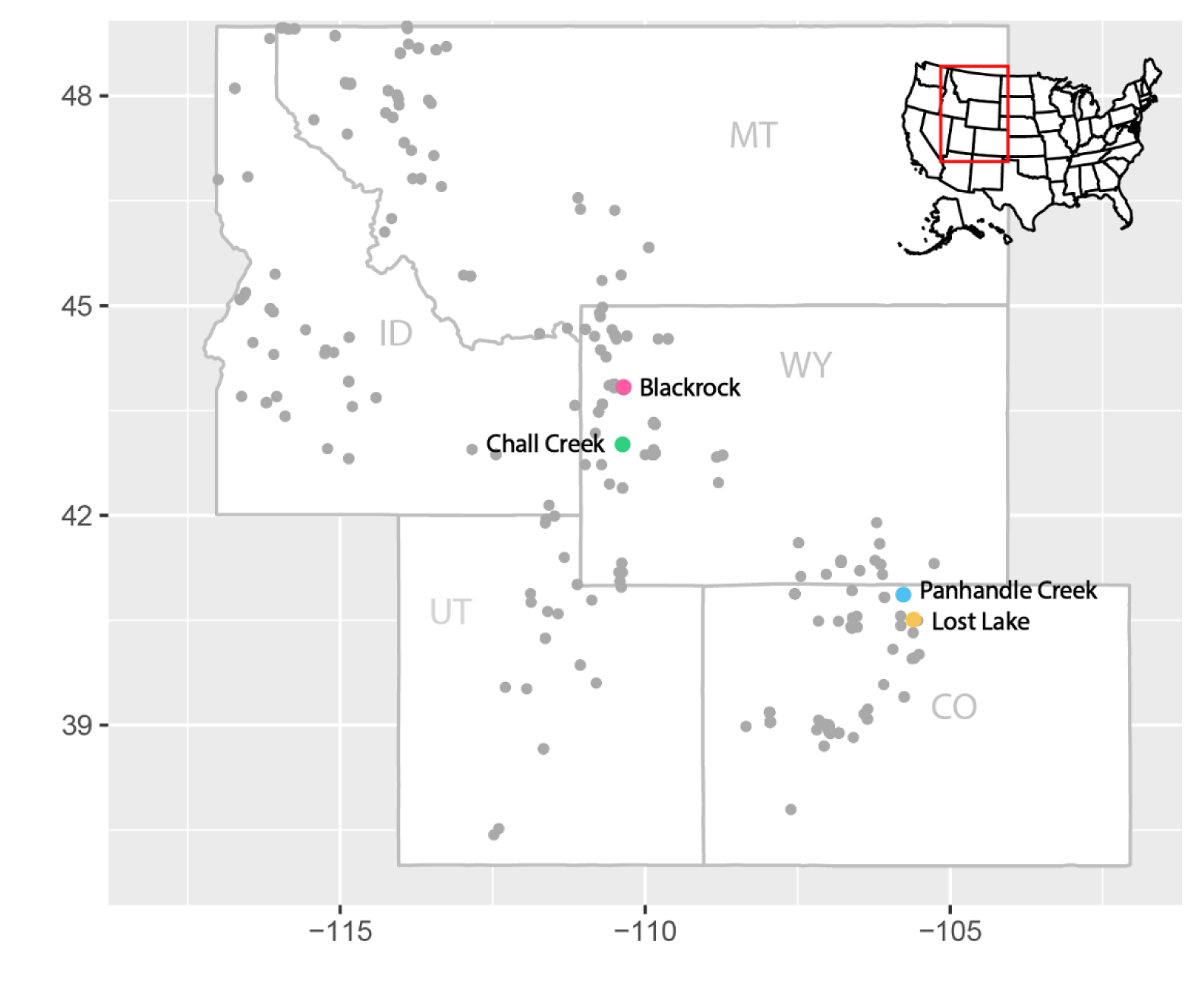
Map of boreal toad (*Anaxyrus boreas boreas*) historical range throughout the Intermountain West, USA. Grey points represent historical and contemporary boreal toad populations. Colored points indicate boreal toad egg mass sources for the experimental exposure study.

### Bd isolation, toad exposure, and monitoring

We isolated Bd from American bullfrog (*Lithobates catesbeianus*) tadpoles collected in Boulder County, CO following Fisher et al. (2018). This isolate had three passages prior to host exposure, limiting potential for pathogen attenuation as documented in other studies (Brem et al. 2013; Langhammer et al. 2013). Recent data also suggest that Bd isolates in Colorado and Wyoming are phylogenetically similar and unlikely to display differences in pathogenicity (Rothstein et al. unpubl. data). To prepare Bd inocula, we grew Bd in sterile 1% Tryptone broth without antibiotics at room temperature for 5-7 days prior to plating on sterile Tryptone-agar media. Plates were incubated at room temperature for another 5-7 days prior to zoospore harvest. We washed plates with 1% Tryptone to gather zoospores while leaving zoosporangia on the plate. This process ensured accurate counts of zoospores with a hemacytometer without biasing counts with zoosporangia. We used a 3×4 factorial design to expose boreal toads from each of the four populations to the three doses of Bd (Table 1). Hemacytometer counts of zoospores informed the dilutions and makeup of the three doses (i.e., 10^2^, 10^3^, 10^5^). In the experimental room, toads were individually exposed to Bd with a 24hr bath in a 15ml solution of Bd in a 150mm x 25mm petri-dish per their treatment dose (Table 1). After 24hrs, toads were placed back into their individual habitat. We checked toads daily to feed, water, and detect mortalities. Beginning seven days post-exposure, and continuing once every seven days until day 56, toads were weighed and swabbed for Bd. We swabbed toads five times on each body surface where Bd is typically found (venter, hands, feet), and did this identically each swab sampling. While swabbing may remove some active Bd zoospores from the host epidermis each week, Kumar et al. (2020) showed that swabbing does not alter fungal loads in an amphibian infected with a similar pathogen, *Batrachochytrium salamandrivorans* (Bsal). Therefore swabbing is unlikely to disturb zoospores that occur deeper in the stratum corneum and stratum granulosum tissue layers (Van Rooij et al. 2015), thus not artificially clearing infections. Swabs were stored in 70% ethanol at −20°C prior to DNA extraction and qPCR (see Appendix 1 for more details). Toads alive on day 56 were humanely euthanized via overdose of MS-222.

**Table 1.** Number of boreal toads (*Anaxyrus boreas boreas*) from each population exposed to each *Batrachochytrium dendrobatidis* (Bd) treatment dose.

| Population | Bd Treatment Dose |  |  |
| --- | --- | --- | --- |
|  | 10<sup>2</sup> (low) | 10<sup>3</sup> (mid) | 10<sup>5</sup> (high) |
| Blackrock, WY | 11 | 12 | 11 |
| Chall Creek, WY | 13 | 14 | 14 |
| Lost Lake, CO | 18 | 21 | 20 |
| Panhandle Creek, CO | 13 | 13 | 13 |

### Data analysis approach

We used a multistate mark-recapture approach (Schwarz et al. 2008; Nichols et al. 1992) to investigate differences in resistance or tolerance to Bd infection among boreal toad populations. This modeling framework allows host weekly survival and state transition probabilities to be modeled as a function of static and dynamic covariates. Static covariates are typical in epidemiological studies and include factors like geographic origin, exposure dose, or host mass prior to exposure. Dynamic covariates are often not incorporated because they change throughout the experiment, such as individual pathogen burden or the change in host mass. Therefore, we modeled weekly survival not only as a function of static covariates (e.g., exposure dose), but also as a function of dynamic individual host (e.g., mass change) and pathogen covariates (e.g., Bd load). This allowed us to capture individual variation and ask questions about which static or dynamic covariates are most important in determining host fitness (i.e., survival). Importantly, this approach allowed us to assess differences in host tolerance by comparing estimates of weekly survival probabilities for different populations across a gradient of pathogen burdens (Figure 1A). Because some toads lost and re-gained Bd infections multiple times throughout the study, a common occurrence in host-pathogen dynamics that is rarely modeled, we were able to investigate the effect of several static or dynamic covariates on the probability an individual transitions from infected to cleared (Ψ^IC^) and cleared to re-infected (Ψ^CI^). These state transition probabilities also serve as lines of evidence for disease resistance (Figure 1D and 1E). Pathogen clearance (Ψ^IC^) implies a direct host mechanism of resisting pathogen growth and presence, and pathogen reinfection implies a lack of resistance to infection (Ψ^CI^). Together, these state transition parameters provide valuable information on host-pathogen dynamics and host resistance relative to simply reporting the raw counts of each transition or ignoring the clearance phenomenon completely.

### Covariates and Hypotheses

We constructed ‘capture’ histories for each individual to estimate weekly survival and state transition probabilities. These histories denote whether each individual is alive and infected (I) or cleared (C), or dead (0) at each weekly sampling period. Using these states, we modeled weekly survival and state transitions as a function of several static or dynamic covariates. Static covariates included an individual’s geographic population of origin (*POP*), geographic region (*REG*), and exposure treatment (*TRT*; Table 2).We also included several individual, time-varying (dynamic) covariates including an individual’s mass (*MASS*), a metric that represented the relative change in body mass (Δ*MASS*; Table 2), and a quantitative index of weekly pathogen burden (Bd load; *BD*). We developed this index of pathogen burden by taking the natural log of the mean PCR copy number + 1 for each sample to use as a quantitative covariate in our analyses and refer to this metric as ‘Bd load’ throughout. These dynamic covariates allowed us to investigate their effects on weekly host survival probabilities, overcoming limitations of classical approaches that only include static, group-level comparisons that describe conditions at the onset of the study.

**Table 2.**
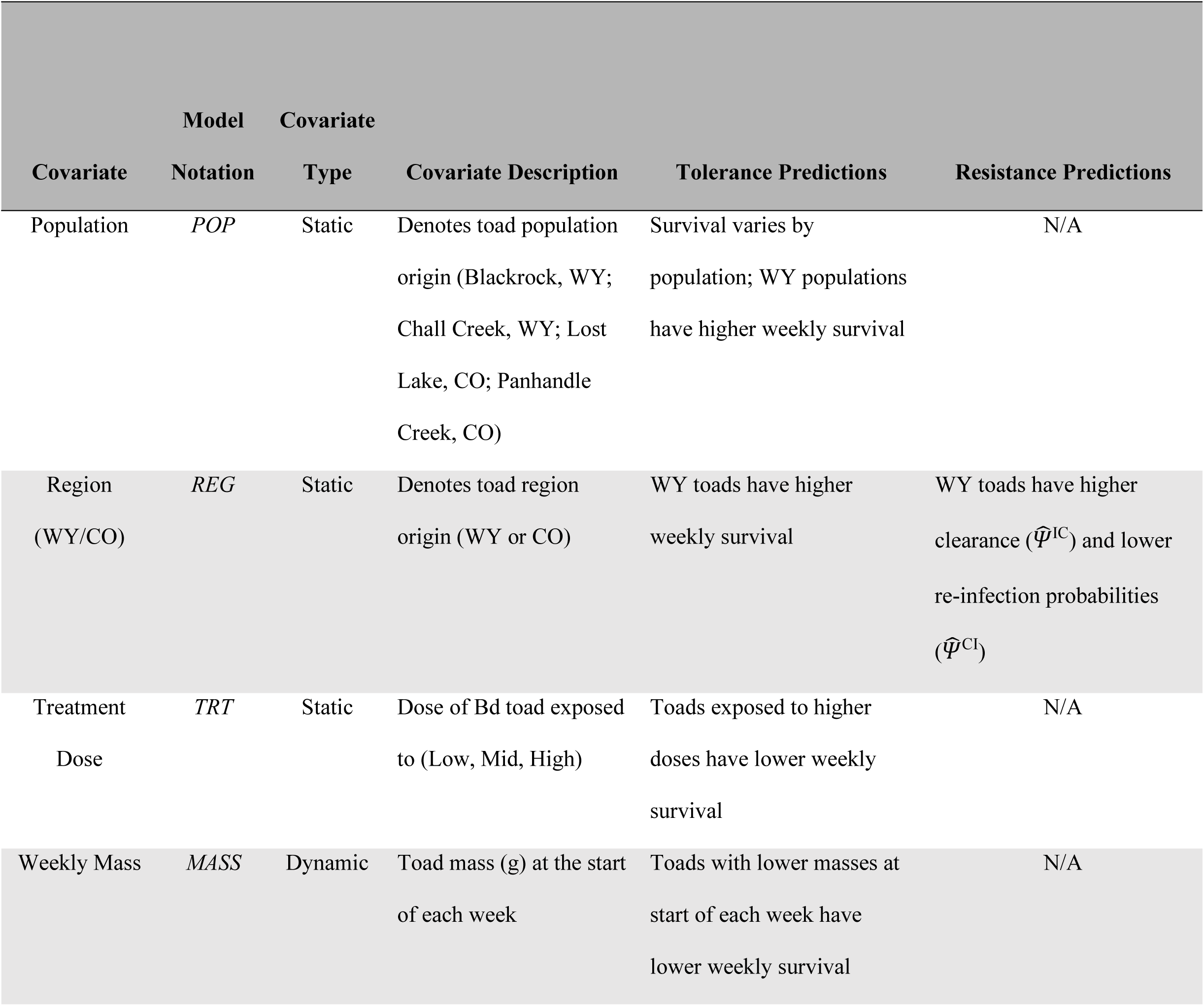
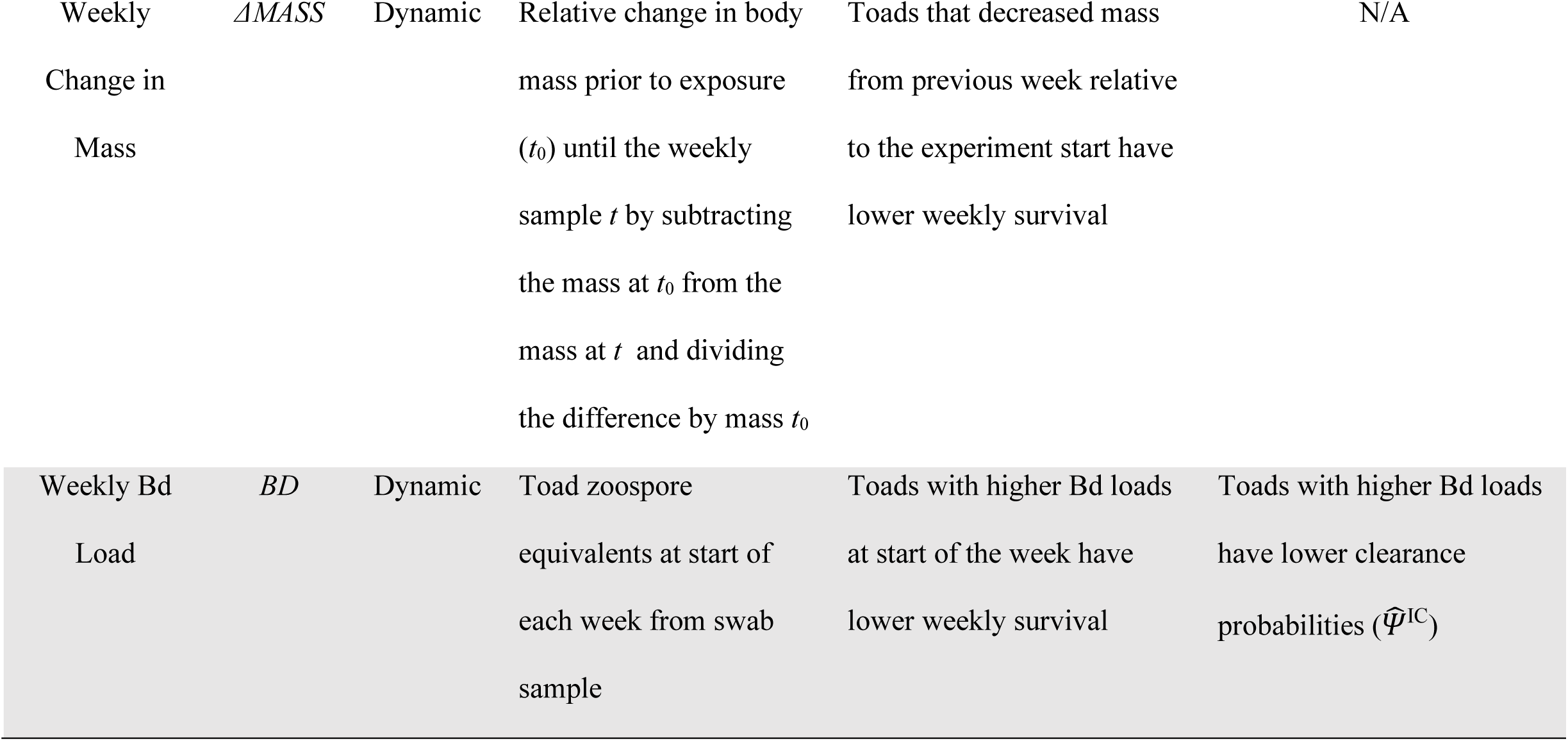
Predicted relationships between multistate model covariates and boreal toad (*Anaxyrus boreas boreas*) tolerance or resistance to *Batrachochytrium dendrobatidis* (Bd). N/A indicates that a given covariate was not applicable to test a specific prediction.

We tested for boreal toad tolerance to Bd by fitting models that reflected differences in weekly survival probability (*S*) attributed to geography (*POP* or *REG*), disease status (*BD* or *TRT*), and body mass (*MASS* or Δ*MASS*) and additive or interactive combinations of these covariates (Table 2). We hypothesized that Wyoming toads would have higher weekly survival per a given Bd load than Colorado toads, either without variation among populations within a region (*REG*) or with variation among populations within and across regions (*POP;* Figure 1A). We also expected that toads with lower Bd would have higher weekly survival, and that this relationship would be better modeled by the individual, time-varying measure of Bd load (*BD*) than by the static factor of initial exposure dose treatment (*TRT*). Finally, we hypothesized that larger toads (*MASS*) or those that had the smallest relative change in body mass (Δ*MASS*) would have higher weekly survival. We also fit several null weekly survival models that reflected the effect of infection alone based on disease state (cleared or infected) or constant survival probabilities for all individuals regardless of disease state, geography, disease status, or body mass.

We tested for boreal toad resistance to Bd in several ways. Using the multistate framework, we hypothesized that if Wyoming toads are more resistant, they would have higher probabilities of clearing infections (Ψ^IC^, Figure 1D; Table 2) and lower probabilities of regaining infections (Ψ^CI^ Figure 1E; Table 2). Therefore, we fit models where these transition probabilities are associated with an individual’s region (*REG*). We did not have enough transitions between states to fit models with population effects (*POP*). We also hypothesized that an individual’s pathogen burden may influence their ability to clear infections between sampling occasions, thus we fit models where this transition probability (Ψ^IC^) was a function of the individual’s Bd load (BD) at time *t*. We also included models where the state transition probabilities (Ψ^IC^ or Ψ^CI^) remained constant from week to week (.).

Additionally, we assessed boreal toad resistance to Bd by testing for differences in the average peak Bd loads (Figure 1B) and average time to reach peak Bd loads (Figure 1C) between Colorado and Wyoming toads. We used t-tests to assess these differences and evaluate the strength of effects by reporting the mean differences and associated measures of precision (95% confidence intervals). If boreal toad populations in Wyoming are more resistant to Bd than those in Colorado, we predicted that Wyoming populations would experience lower peak Bd loads, on average, or take longer to reach peak loads.

### Model building and fitting

We fit all models using the multistate model type in *Program MARK* (White and Burnham 1999). We employed a secondary candidate set model building approach (Bromaghin et al. 2013), where we fit models to test hypotheses about one focal parameter at a time (i.e., *S*) using a null model structure for the other, non-focal parameters (i.e., Ψ^IC^ or Ψ^CI^, Appendices 2-3). This approach is better than other model building strategies at recovering total Akaike weights, limiting inclusion of unimportant covariates, and identifying the top model when using model types with multiple parameters (Morin et al. 2020). Our final ‘combined model set’ included all combinations of supported structures (with AICc < 10) from each of the focal parameter model sets (i.e., weekly survival and state transition probabilities).

## RESULTS

To address hypotheses regarding boreal toad tolerance and factors influencing host weekly survival (*S*), our focal model set contained 49 survival models (Appendix 2), two of which were well-supported (ΔAIC_C_ < 10) and retained in our final combined model set. To test hypotheses regarding boreal toad resistance and factors influencing host weekly transition probabilities (Ψ^IC^ and Ψ^CI^) we fit six transition probability models (Appendix 3) and all were supported (ΔAIC_C_ < 10) and retained in our final candidate model set. Our final candidate model set consisted of 12 models that included all combinations of the six state transition probability structures and two survival structures (Table 3).

**Table 3.**
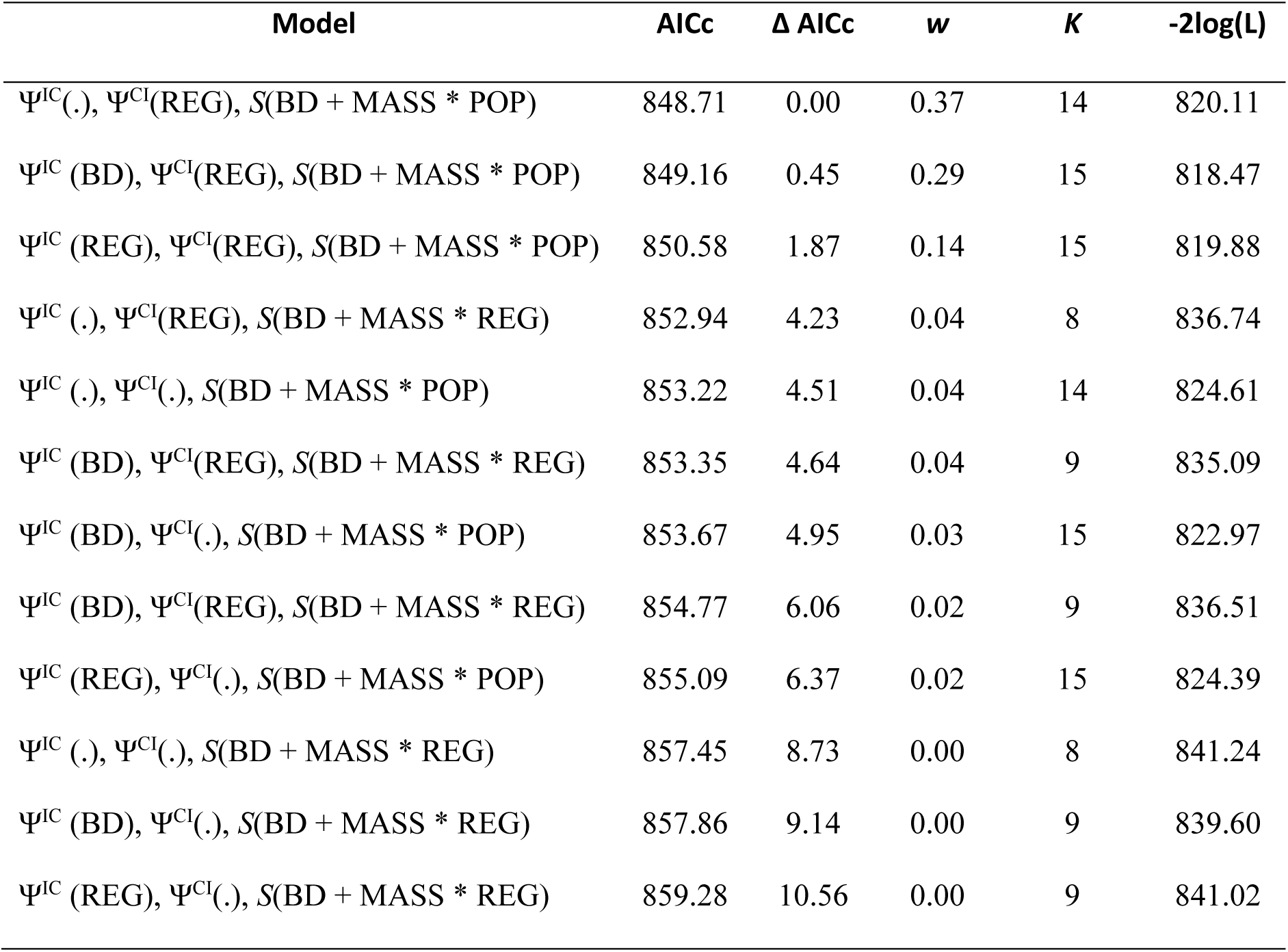
Model selection results from multistate mark-recapture models of boreal toads (*Anaxyrus boreas boreas*) exposed to *Batrachochytrium dendrobatidis* (Bd) in a laboratory experiment. Parameters include: Ψ^IC^, transition probability from infected to cleared; Ψ^CI^, transition probability from cleared to infected; *S*, weekly survival probability. Covariates include REG, regional variation in host source (Colorado or Wyoming); POP, population variation in host source; BD, weekly Bd load; MASS, weekly measurement of mass in grams; (.), no variation.

| Model | AICc | $\Delta$ AICc | $w$ | $K$ | -2log(L) |
| --- | --- | --- | --- | --- | --- |
| $\Psi^{IC}(.), \Psi^{CI}(\text{REG}), S(\text{BD} + \text{MASS} * \text{POP})$ | 848.71 | 0.00 | 0.37 | 14 | 820.11 |
| $\Psi^{IC}(\text{BD}), \Psi^{CI}(\text{REG}), S(\text{BD} + \text{MASS} * \text{POP})$ | 849.16 | 0.45 | 0.29 | 15 | 818.47 |
| $\Psi^{IC}(\text{REG}), \Psi^{CI}(\text{REG}), S(\text{BD} + \text{MASS} * \text{POP})$ | 850.58 | 1.87 | 0.14 | 15 | 819.88 |
| $\Psi^{IC}(.), \Psi^{CI}(\text{REG}), S(\text{BD} + \text{MASS} * \text{REG})$ | 852.94 | 4.23 | 0.04 | 8 | 836.74 |
| $\Psi^{IC}(.), \Psi^{CI}(.), S(\text{BD} + \text{MASS} * \text{POP})$ | 853.22 | 4.51 | 0.04 | 14 | 824.61 |
| $\Psi^{IC}(\text{BD}), \Psi^{CI}(\text{REG}), S(\text{BD} + \text{MASS} * \text{REG})$ | 853.35 | 4.64 | 0.04 | 9 | 835.09 |
| $\Psi^{IC}(\text{BD}), \Psi^{CI}(.), S(\text{BD} + \text{MASS} * \text{POP})$ | 853.67 | 4.95 | 0.03 | 15 | 822.97 |
| $\Psi^{IC}(\text{BD}), \Psi^{CI}(\text{REG}), S(\text{BD} + \text{MASS} * \text{REG})$ | 854.77 | 6.06 | 0.02 | 9 | 836.51 |
| $\Psi^{IC}(\text{REG}), \Psi^{CI}(.), S(\text{BD} + \text{MASS} * \text{POP})$ | 855.09 | 6.37 | 0.02 | 15 | 824.39 |
| $\Psi^{IC}(.), \Psi^{CI}(.), S(\text{BD} + \text{MASS} * \text{REG})$ | 857.45 | 8.73 | 0.00 | 8 | 841.24 |
| $\Psi^{IC}(\text{BD}), \Psi^{CI}(.), S(\text{BD} + \text{MASS} * \text{REG})$ | 857.86 | 9.14 | 0.00 | 9 | 839.60 |
| $\Psi^{IC}(\text{REG}), \Psi^{CI}(.), S(\text{BD} + \text{MASS} * \text{REG})$ | 859.28 | 10.56 | 0.00 | 9 | 841.02 |

### Tolerance

Overall, we found strong support for intraspecific variation in Bd tolerance among boreal toad populations. Boreal toads from populations in Wyoming consistently displayed higher weekly survival probabilities at a given Bd load when compared to those in Colorado, particularly as Bd loads increased (Figure 3). Toads from populations in Colorado were also more variable. Panhandle Creek toads exhibited moderate weekly survival compared to those from Wyoming, while Lost Lake toads exhibited the lowest weekly survival especially at higher Bd loads. This effect of Bd load was mediated by an interaction between population origin and weekly body mass. For example, if comparing weekly survival of toads that weighed 0.4 grams across populations at an identical Bd load of 12, those in Colorado had lower weekly survival probabilities (Lost Lake = 0.43, 95% CI [0.31-0.56]; Panhandle Creek = 0.50, 95% CI [0.36-0.65]; Figure 4) compared to Wyoming toads (Blackrock = 0.71, 95% CI [0.58-0.82]; Chall Creek = 0.81, 95% CI [0.68-0.90]; Figure 4).

**Figure 3.**
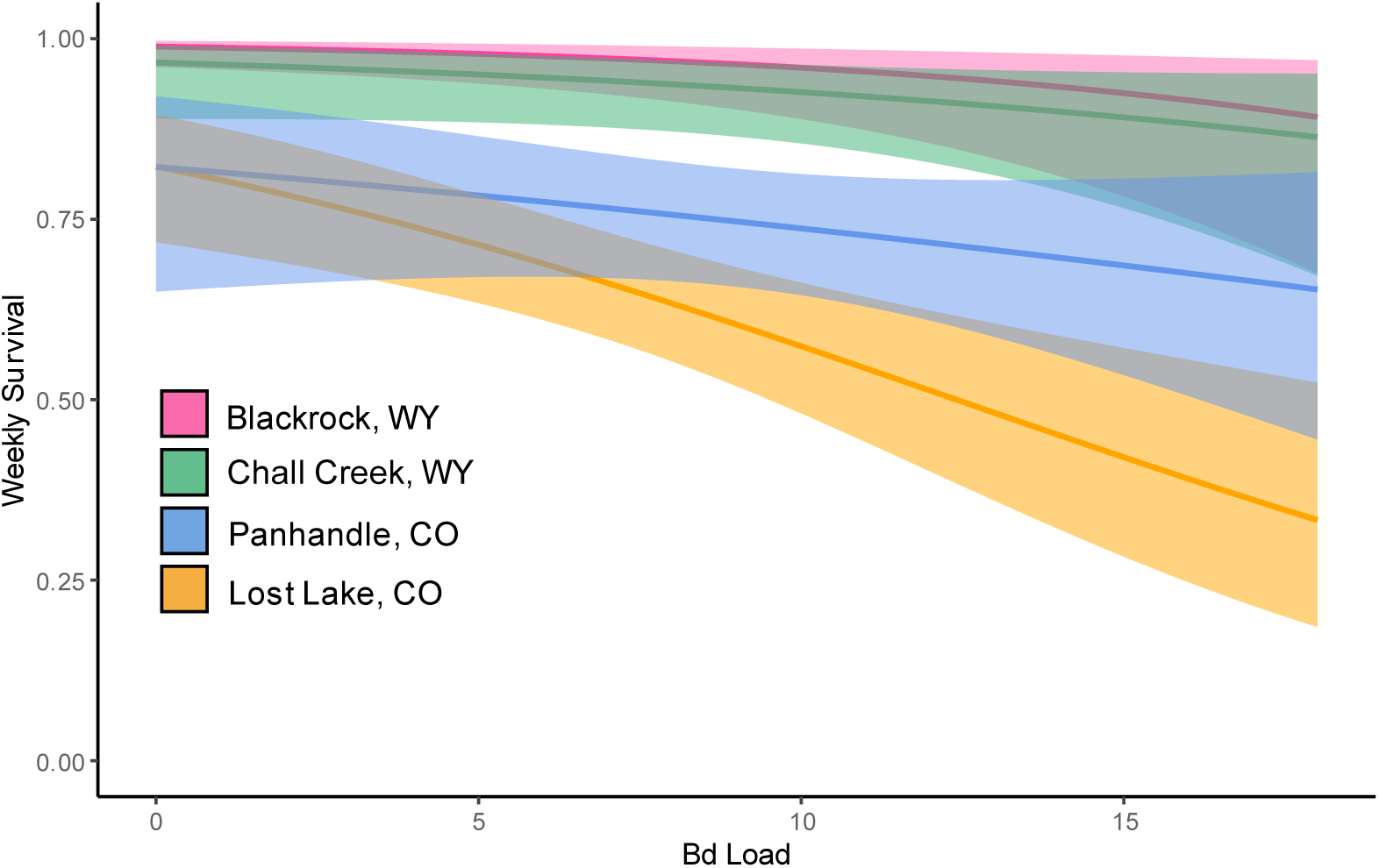
Estimated weekly survival probabilities of boreal toads (*Anaxyrus boreas boreas*) from four populations exposed to *Batrachochytrium dendrobatidis* (Bd). Survival estimates (solid lines) vary as a function of Bd load, ln(mean copy number +1). Estimates are based on the top model, *S*(BD + MASS * POP), and reported for toad mass = 0.50 g. Bands represent 95% confidence intervals.

**Figure 4.**
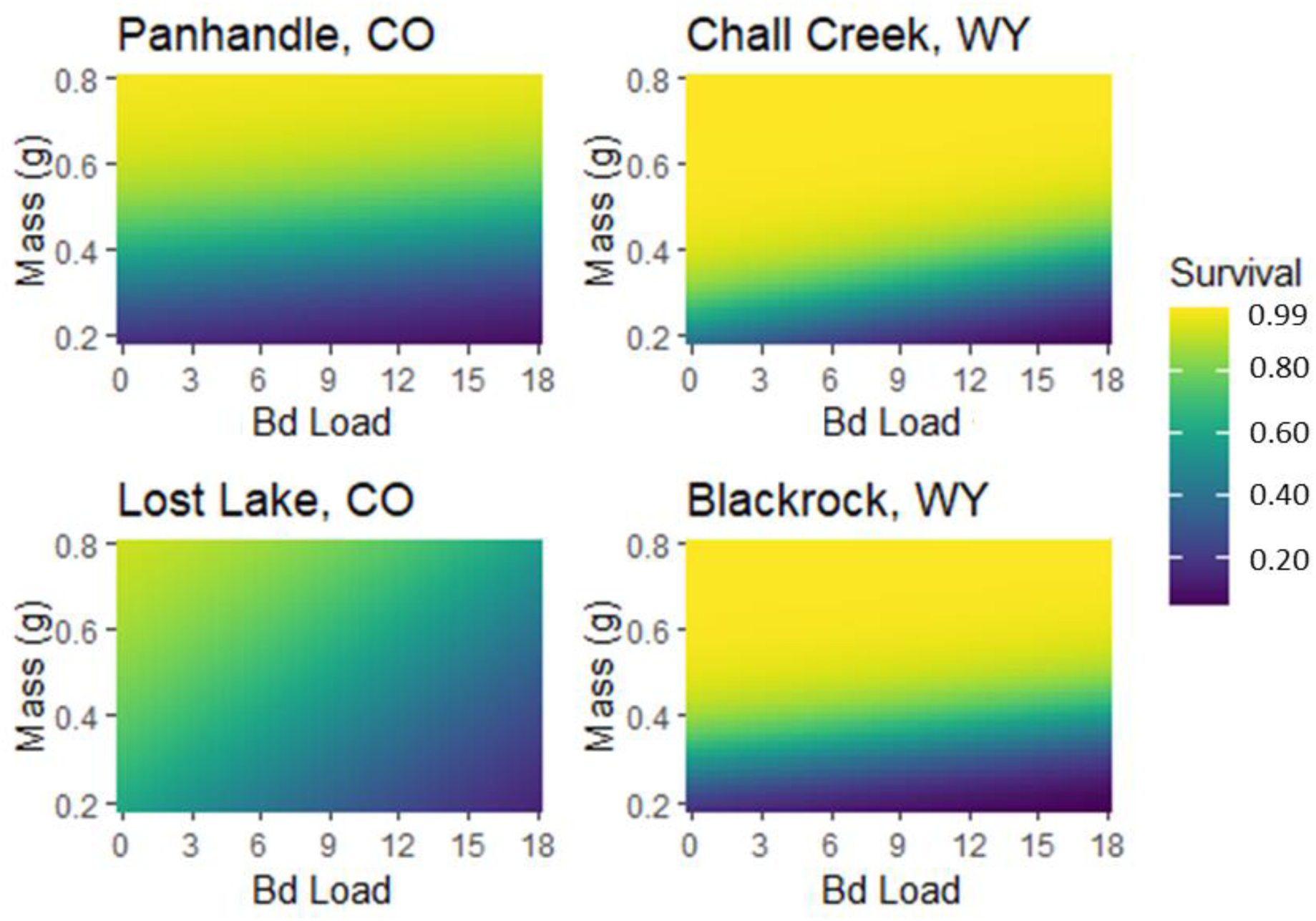
Heatmaps displaying the interaction between mass (g) and *Batrachochytrium dendrobatidis* (Bd) load on weekly survival probability for four populations of boreal toads (*Anaxyrus boreas boreas*). Yellow and green colors indicate high weekly survival estimates where darker blues and purples represent lower weekly survival estimates. The range in mass (g) is condensed to 0.20 - 0.80g to better show the gradient of survival estimates. Individuals ranged from 0.20 - 1.12g during the experiment, but only six individuals from Wyoming populations weighed more than 0.80g.

### Resistance

We found mixed support for increased resistance of Wyoming boreal toads to Bd compared to those in Colorado. The probability an infected toad cleared Bd was low (Ψ̂^IC^ =0.05) and similar between regions (Figure 5). However, Wyoming boreal toads that did clear Bd infections had a much lower probability of being reinfected (Ψ̂^CI^ = 0.16; 95% CI [0.07, 0.32]) compared to those from Colorado (Ψ̂^CI^ = 0.50; 95% CI [0.22, 0.78]; Figure 5). We also observed differences in weekly survival probabilities among toads that completely cleared their infections (see y-intercept of Figure 3).

**Figure 5.**
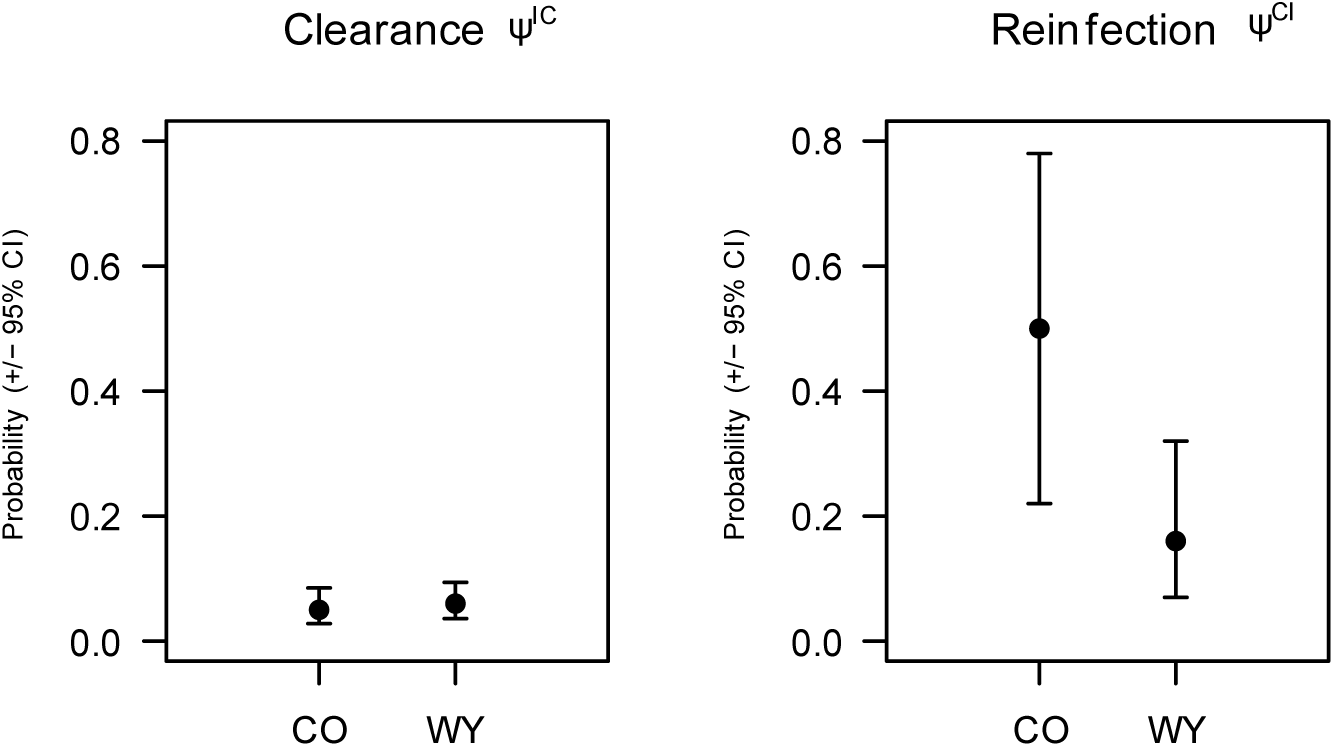
Estimated state transition probabilities (ψ) of boreal toads (*Anaxyrus boreas boreas*) from Colorado (CO) and Wyoming (WY) exposed to *Batrachochytrium dendrobatidis* (Bd). Colorado toads have an equal probability of Bd clearance, but higher probability of becoming reinfected than WY toads.

We found no evidence that average peak Bd loads differed for toads from Colorado and Wyoming (t = −1.30, p-value=0.19, mean difference = −0.62; 95% CI [-1.57, 0.32]; Figure 6). However, boreal toads from Wyoming reached peak Bd loads later than Colorado toads (t = - 2.10, p=0.04, mean difference = −4.24 days; 95% CI [-8.26, −0.22]; Figure 7).

**Figure 6.**
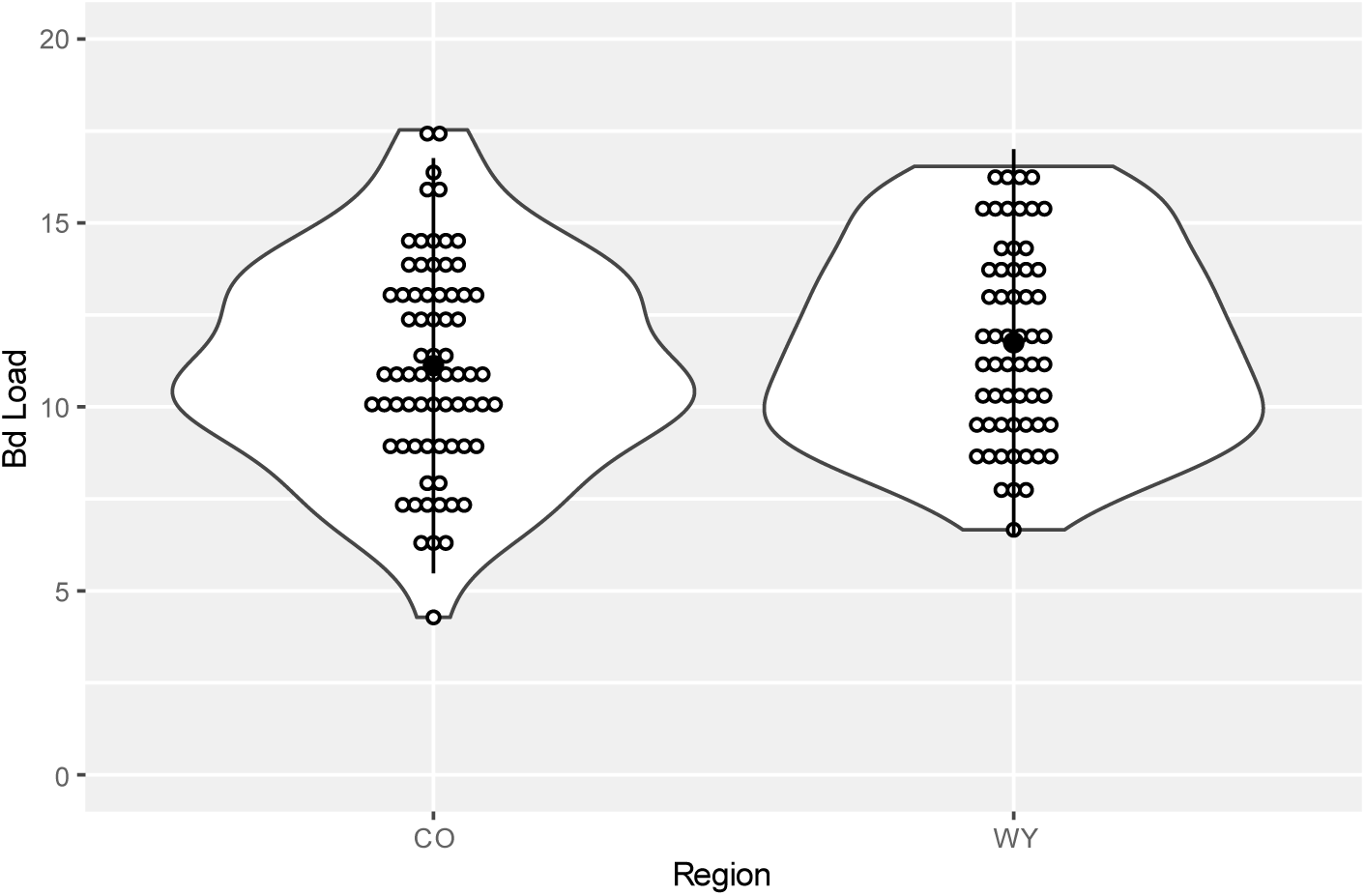
Peak pathogen (*Batrachochytrium dendrobatidis*) loads (Bd load) experienced by exposed boreal toads (*Anaxyrus boreas boreas*) from Colorado (CO) and Wyoming (WY) are similar. Black filled circles denote the region-specific mean, open circles represent individual toads. Vertical lines indicate ± 1 standard deviation from the mean.

**Figure 7.**
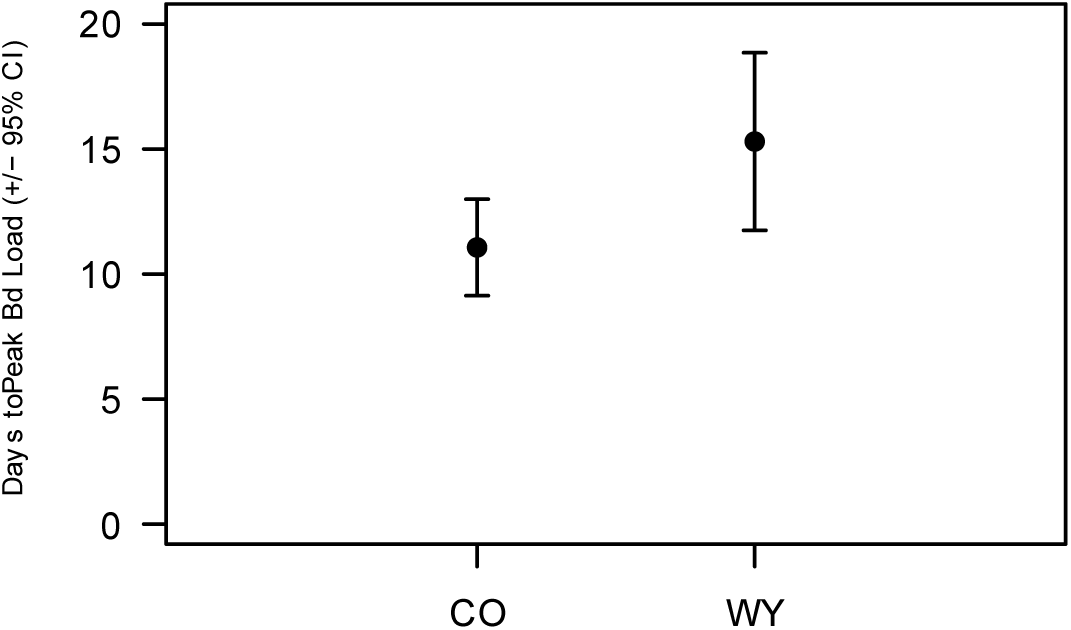
The average number of days for boreal toads (*Anaxyrus boreas boreas*) from Colorado (CO) to reach peak *Batrachochytrium dendrobatidis* (Bd) loads are lower than those from Wyoming (WY).

## DISCUSSION

Quantifying intraspecific variation in host susceptibility to pathogens and parasites is essential to better understand host-pathogen dynamics and inform host conservation efforts. Using boreal toads and Bd as an example of a widespread and highly variable host-pathogen system, we uncovered hidden disease dynamics with a multistate modelling approach. Our results revealed intraspecific variation in tolerance and resistance among host populations and provide targeted guidance for ongoing conservation efforts.

We documented differences in disease tolerance across replicate boreal toad populations in Wyoming and Colorado. Per Bd load, boreal toads from Wyoming had higher weekly survival estimates on average than those from Colorado. The exact mechanisms of how boreal toads from Wyoming are tolerating Bd infections relatively better is still unknown. Tolerance mechanisms are, in general, much more understudied than resistance mechanisms (Råberg et al. 2009) and typically exist at the cellular/tissue/organ scales that work to limit damage caused by both the pathogen and the host immune response and to maintain homeostasis (Medzhitov et al. 2012). Host tolerance mechanisms associated with Bd infection are hypothesized to include stress and damage responses, tissue repair, and factors that limit immunopathology (Grogan et al. 2023). Recent research investigating the differential expression of genes in Bd-infected boreal toads may shed some light on the mechanisms of tolerance (Corey-Rivas et al. *pers. comm.*). Corey-Rivas et al. found that Bd-infected boreal toads from Utah had upregulated immune responses, and Colorado boreal toads had downregulated molecular functioning, suggesting a disruption of electrolyte balance. Importantly, sampled Utah populations may be more closely related to the boreal toad populations in western Wyoming (Goebel et al. 2009; Oyler-McCance et al. 2017). This promising research may begin to link which gene regions are associated with Bd tolerance and provide metabolic markers to aid future genetic studies. Ultimately these findings, along with our own, could guide genomic supplementation or captive breeding strategies to enhance the prevalence of certain tolerant genotypes to aid species persistence on the landscape.

We also observed some evidence for Wyoming boreal toad resistance to Bd. Resistance mechanisms of amphibian hosts to Bd are well-studied and include functional immune responses (Grogan et al. 2018; Rollins-Smith 2001), anti-microbial peptides (Woodhams et al. 2006; Rollins-Smith and Conlon 2005; Rollins-Smith 2009), anti-fungal skin flora (Rebollar et al. 2020), or other physiological traits (e.g., increased skin sloughing; Ohmer et al. 2017). Behavioral strategies that promote amphibian disease resistance are particularly intriguing. For example, researchers studying the effects of a related pathogenic fungus (*Batrachochytrium salamandrivorans*; Bsal) on newts (*Ichthyosaura alpestris*) found that infected individuals routinely switched habitats from aquatic to terrestrial that aided the desiccation of the fungus and clearing their infections (i.e., resistance; Daversa et al. 2018). In fact, a field study of radio-telemetered boreal toads at a different population in western Wyoming from one of ours sampled, found that infected individuals preferentially select open habitats that are warmer and have higher probabilities of lowering their Bd burden through behavioral thermoregulation (Barrile et al. 2021). We did not witness individual toads moving between aquatic or terrestrial habitats within their individual containers, but future studies could include video recording of several individuals in each treatment. Behavioral differences may have the potential to explain the lower probabilities of becoming reinfected and the slightly longer times to reach peak pathogen burdens in Wyoming boreal toads that we observed.

In addition to differences in tolerance and resistance observed between Wyoming and Colorado boreal toads, weekly survival estimates also revealed differences among regions in the absence of Bd (y-intercept Figure 2). Colorado toads that cleared their infections (Bd load = 0) had weekly survival probabilities that were lower (∼12% difference) than boreal toads from Wyoming who had also cleared their infections. In other words, uninfected Colorado boreal toads have lower weekly survival probabilities than uninfected Wyoming toads. This reduced fitness in Colorado toads that cleared previous infections could indicate a lag effect from their prior infection and the cost of clearing their infection ultimately resulted in lower survival probabilities. Alternatively, this difference could imply that Colorado toads have inherently lower vigor, attributed to the lower genomic diversity observed in Colorado populations (Trumbo et al. *In Revision*) that could potentially contribute to increased incidence of inbreeding depression (Keller and Waller 2002).

Our findings of decreased tolerance and resistance to Bd in Colorado populations of boreal toads and the possibility of lower innate vigor, may stem from their phylogeographic history and contemporary population genetics. Overall genomic diversity is the source for genetic adaptation and is predictive of both inbreeding depression and adaptive potential (Kardos et al. 2021). Therefore, we might expect that boreal toad populations in western Wyoming possess greater variation in beneficial Major Histocompatibility Complex (MHC) genotypes for natural selection to act on than those in Colorado. Previous research on amphibian susceptibility to Bd has found that some species have MHC profiles and subsequent immune responses that improve Bd resistance (Savage and Zamudio 2011; Bataille et al. 2015; Savage et al. 2018; Trujillo et al. 2021). In fact, research outside of the amphibian-Bd system has shown that specific MHC genotypes are correlated with unique microbial communities in mice (Kubinak et al. 2015), highlighting the potential for MHC diversity to influence anti-fungal microbial diversity on amphibian skin, a known resistance mechanism (Harris et al. 2006; Rebollar et al. 2020). Further research on the MHC diversity of boreal toads across regions is warranted to investigate whether certain MHC genotypes exist in specific populations, and if those genotypes contribute to Bd resistance.

Furthermore, the boreal toad populations in our study area are currently disjunct and may have experienced restricted gene flow for a long time, owing to the unsuitable habitat and low elevations between them (e.g., the Red Desert of Wyoming; Hammerson et al. 1999) and the likely colonization of Colorado after glacial retreat during the last glacial maximum (∼12kya; Brugger et al. 2019; Guido et al. 2007). There is evidence of strong population genetic divergence between these regions using mtDNA markers (Goebel et al. 2009), but mixed evidence among a comparison of different genetic and genomic marker types (Oyler-McCance et al. 2017). Regardless, boreal toad populations in Colorado have low genomic diversity (Trumbo et al. *Accepted*), potentially influencing a lack of adaptive response to a novel stressor (i.e., disease) over the last 50 years as a function of high inbreeding depression. In contrast, boreal toad populations in northwestern Wyoming have higher genetic variation than those in Colorado (Oyler-McCance et al. 2017, Trumbo et al. *Accepted*). Thus, the potential for historical isolation between these two regions may have contributed to the low genomic diversity found in boreal toad populations in Colorado, where populations exist at their elevational limits and are now fragmented across the landscape due primarily to Bd-related local extirpations (Mosher et al. 2018; Trumbo et al. *Accepted*).

Our findings have significant implications for management of boreal toads in the southern Rocky Mountains. Bd is widespread and highly persistent in our study area (Mosher et al. 2018), supporting the idea that tolerance mechanisms would be a better host strategy over resistance. Given our findings of a strong signal of tolerance in Wyoming boreal toads over weak evidence for resistance, conservation measures that attempt to eradicate Bd on the landscape are likely not optimal (Gerber et al. 2023). With the knowledge that boreal toad genomic diversity is low in the southern Rocky Mountain region (Trumbo et al. *Accepted*) and with new evidence that tolerance and resistance to Bd is also low (this study), success of reintroduction efforts may be improved by incorporating genetic material from other sources. Reintroduction and translocation efforts to conserve and recover boreal toads in Colorado currently follow a nearest-neighbor approach where source populations are chosen based on distance to the selected reintroduction site (H Crockett, Colorado Parks and Wildlife, pers. comm.). Indeed, high-profile success stories from the Florida panther (*Puma concolor*) illustrate the benefits of reintroducing and translocating individuals from different genetic lineages to mask deleterious alleles and improve vigor and resiliency (Johnson et al. 2010). In addition to reintroduction and translocation implications, our findings have strong potential to influence captive rearing and breeding efforts. Managers might consider introducing genetic material outside of Colorado and southeastern Wyoming to improve the potential for heritable tolerance traits to be carried in crosses between lineages. Further research should evaluate the heritability of tolerance in boreal toads.

In conclusion, we provide evidence that intraspecific variation in host tolerance and resistance are responsible for shaping regional differences in host-pathogen dynamics in response to a deadly fungal pathogen. We also leverage the power of multistate modeling to reveal traditionally hidden disease dynamics that contribute to our understanding of the disease ecology of our system and we encourage its use in future experimental exposures of pathogens or parasites to hosts. Finally, by documenting differences in host defense strategies across a large portion of a host’s range, we demonstrate that characterizing an entire species as ‘tolerant’ or ‘resistant’ can be unwise and potentially detrimental to host conservation as a whole without accounting for intraspecific variation.

## ACKNOWLEDGMENTS

This project received funding from US Geological Survey, the Rocky Mountain National Park Bailey Fellowship, NSF Graduate Research Fellowship, and a Colorado State University Graduate Degree Program in Ecology small grant. We thank J. Kerby for use of his laboratory and training in qPCR. We thank A. Reed, K. Mauerman, E. Hazzard, and E. Doorack for assisting with animal husbandry and data collection. We thank K. Hoke for microscope use and K. Huyvaert for supplies and advice, in addition to both providing helpful feedback on earlier versions of this manuscript. We thank W. Estes-Zumpf of Wyoming Department of Game and Fish, F.B. Wright III, H. Crockett, and B. Neuschwanger of Colorado Parks and Wildlife for collecting and rearing toads. We thank K. Bestgen for lab troubleshooting and aiding in animal rearing.

## APPENDIX

## Appendix 1 Supplementary Methods Details

### Site selection, egg collection, and animal husbandry

In collaboration with state and federal biologists, we collected up to 250 eggs (i.e., < 5% of available eggs) at each population. Eggs were collected and held in a large Ziploc® (SC Johnson, Racine, Wisconsin, USA) bag filled with water and placed inside cylindrical insulated coolers for transport to a facility for rearing. Eggs were housed at the Colorado Parks and Wildlife (CPW) Fish Research Hatchery in Bellvue, CO within 24h of collection.

Upon arrival at the hatchery, eggs were counted and inspected for viability. Eggs from each population were placed in 50-gallon aquaria and reared together until Gosner stage 30 (Gosner 1960). At this stage, larvae (i.e., tadpoles) were transferred to the Colorado State University Aquatic Research Laboratory and were haphazardly assigned to 10-gallon tanks with ∼ 100 individuals per tank. Fifty percent of the water in aquaria was changed daily with natural well water that ranged from 15-18 °C. Tadpoles were fed algae wafers and mixtures of puréed squash, zucchini, and collard greens daily and experienced a 12hr light-dark cycle. Once tadpoles began metamorphosis and developed limbs, we transferred individuals to transition containers consisting of a small plastic container of shallow water with stones to assist larval transition from aquatic to terrestrial habitat through metamorphosis. Fully terrestrial toadlets were then moved to 10-gallon terraria consisting of Eco Earth® (Zoo Med Laboratories Inc., San Luis Obispo, California, USA) substrate and water dishes. Toads were fed flightless *Drosophila melanogaster* dusted in vitamins *ad libidum*. Terraria housed up to 50 toads from the same collection site.

### Swab DNA extraction and qPCR

We used Qiagen Blood and Tissue DNEasy® spin-column kits and followed the manufacturer’s protocol (Qiagen, Venlo, Netherlands) for animal tissues to extract fungal DNA from swab samples with alterations to the tissue digestion methods. Alterations included keeping the swabs in buffer in Step 1 and transferring swabs to spin columns for the primary centrifuge cycle (Step 5). We performed fast-qPCR of extracted DNA in triplicate following established protocols (Kerby et al. 2013).

## Appendix 2: Transition probability model set

Model selection results for six state transition probability (ψ) structures fit to boreal toad capture history data after exposure to *Batrochochytrium dendrobatidis* (Bd) in a laboratory experiment using a multistate model. Model names, Akaike Information Criterion for small sample sizes (AICc), ΔAICc, model weights (*w*), number of parameters (*K*), and *-2log(L)* are given for each model. We modeled the weekly probability an individual transitions from Bd infected, to cleared (ψ^IC^), as a function of Bd load (BD), geographic region of origin of the host (REG), or as a constant (.). We modeled the weekly probability an individual transitions from cleared, back to infected (ψ^CI^), as a function of the geographic region of origin of the host (REG), or as a constant (.). Because the focal parameters of interest were ψ, we only included a basic structure for weekly survival (*S*), allowing survival to be different for infected (I) versus cleared (C) individuals.

| Model | AICc | $\Delta$ AICc | $w$ | $K$ | $-2\log(L)$ |
| --- | --- | --- | --- | --- | --- |
| $\Psi^{IC}(.), \Psi^{CI}(\text{REG}), S(I/C)$ | 998.60 | 0.00 | 0.35 | 5 | 988.52 |
| $\Psi^{IC}(\text{BD}), \Psi^{CI}(\text{REG}), S(I/C)$ | 999.0 | 0.39 | 0.29 | 6 | 986.88 |
| $\Psi^{IC}(\text{REG}), \Psi^{CI}(\text{REG}), S(I/C)$ | 1000.41 | 1.81 | 0.14 | 6 | 988.29 |
| $\Psi^{IC}(.), \Psi^{CI}(.), S(I/C)$ | 1001.1 | 2.48 | 0.10 | 4 | 993.02 |
| $\Psi^{IC}(\text{BD}), \Psi^{CI}(.), S(I/C)$ | 1001.47 | 2.86 | 0.08 | 5 | 991.38 |
| $\Psi^{IC}(\text{REG}), \Psi^{CI}(.), S(I/C)$ | 1002.88 | 4.28 | 0.04 | 5 | 992.80 |

## Appendix 3: Weekly survival probability model set

Model results for the 49 weekly survival probability (*S*) structures fit to boreal toad capture history data after exposure to *Batrochochytrium dendrobatidis* (Bd) in a laboratory experiment using a multistate model. Model names, Akaike Information Criterion for small sample sizes (AICc), ΔAICc, model weights (*w*), number of parameters (*K*), and *-2log(L)* are given for each model. We modeled weekly survival probability for individual toads as a function of static and/or dynamic covariates of interest and additive or interactive combinations where appropriate. Static covariates include: exposure treatment dose of Bd at the start of the experiment (i.e., low, medium, or high; TRT), host population of origin (POP), or host geographic region of origin (REG). Dynamic covariates include: the weekly measure of Bd load (BD), the weekly measure of body mass (MASS), or the weekly change in body mass from time *t* to time *t_0_* (ΔMASS). Two null models were also fit, one with weekly survival as a constant (.), and one with weekly survival differing between infected (I) and cleared (C) individuals. Because the focal parameter of interest was *S*, we only included constant null structures (.) associated with weekly state transition parameters (ψ).

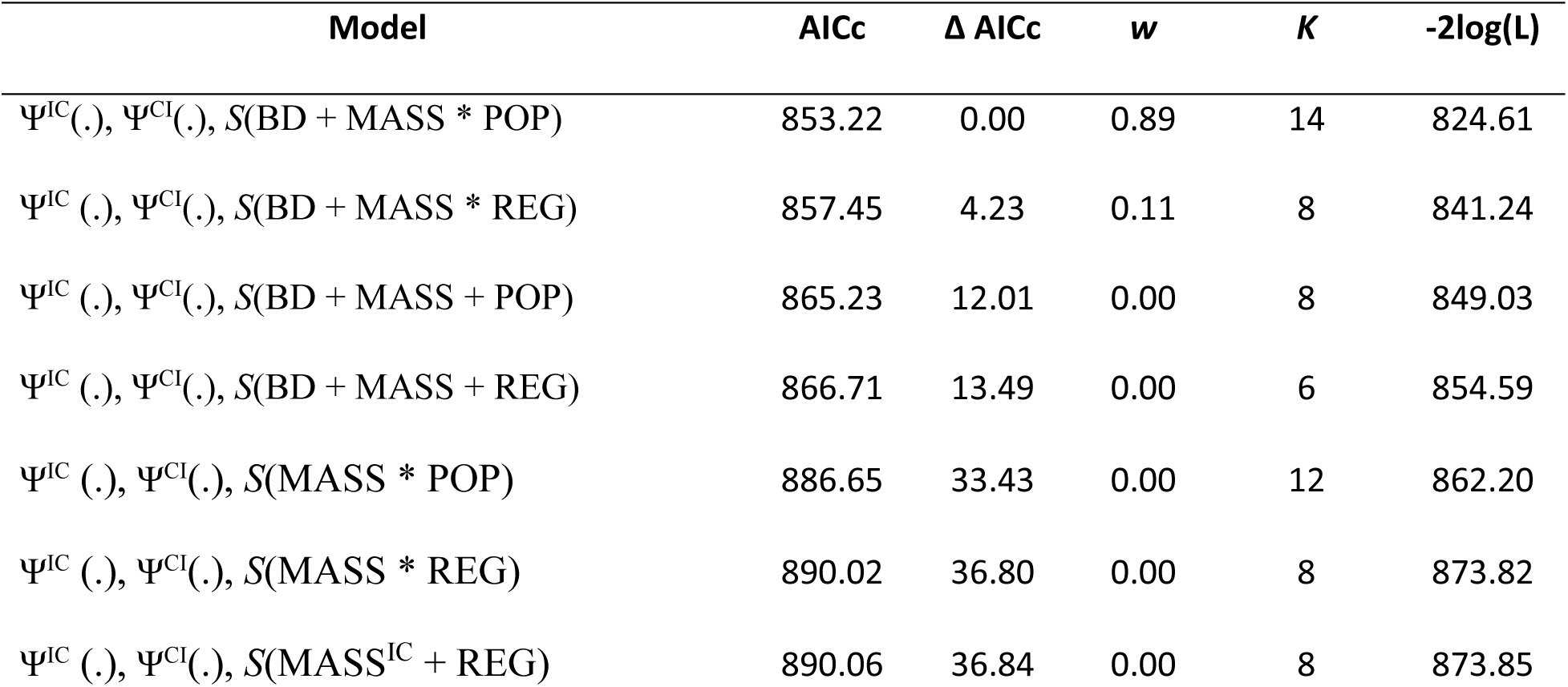
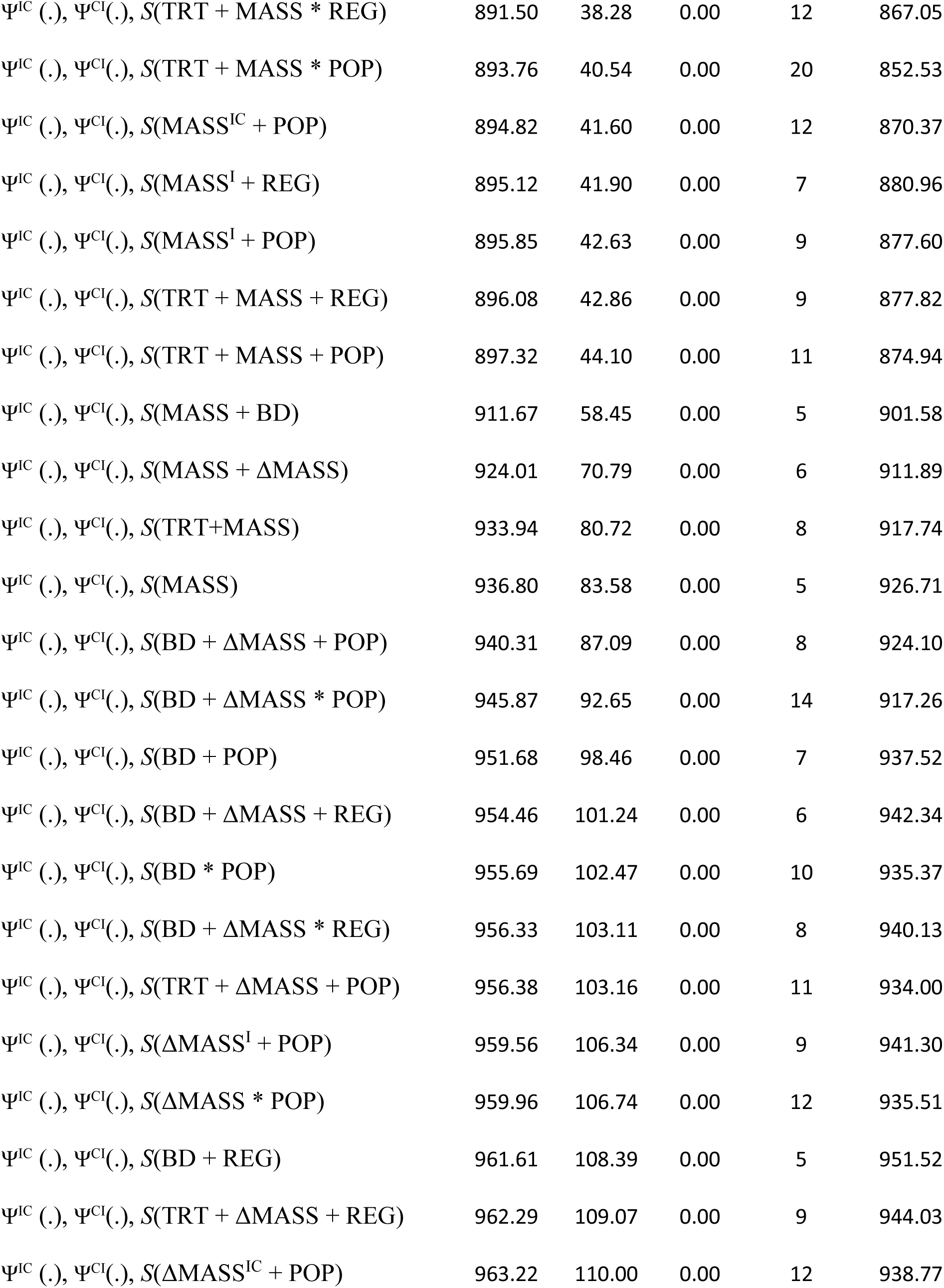
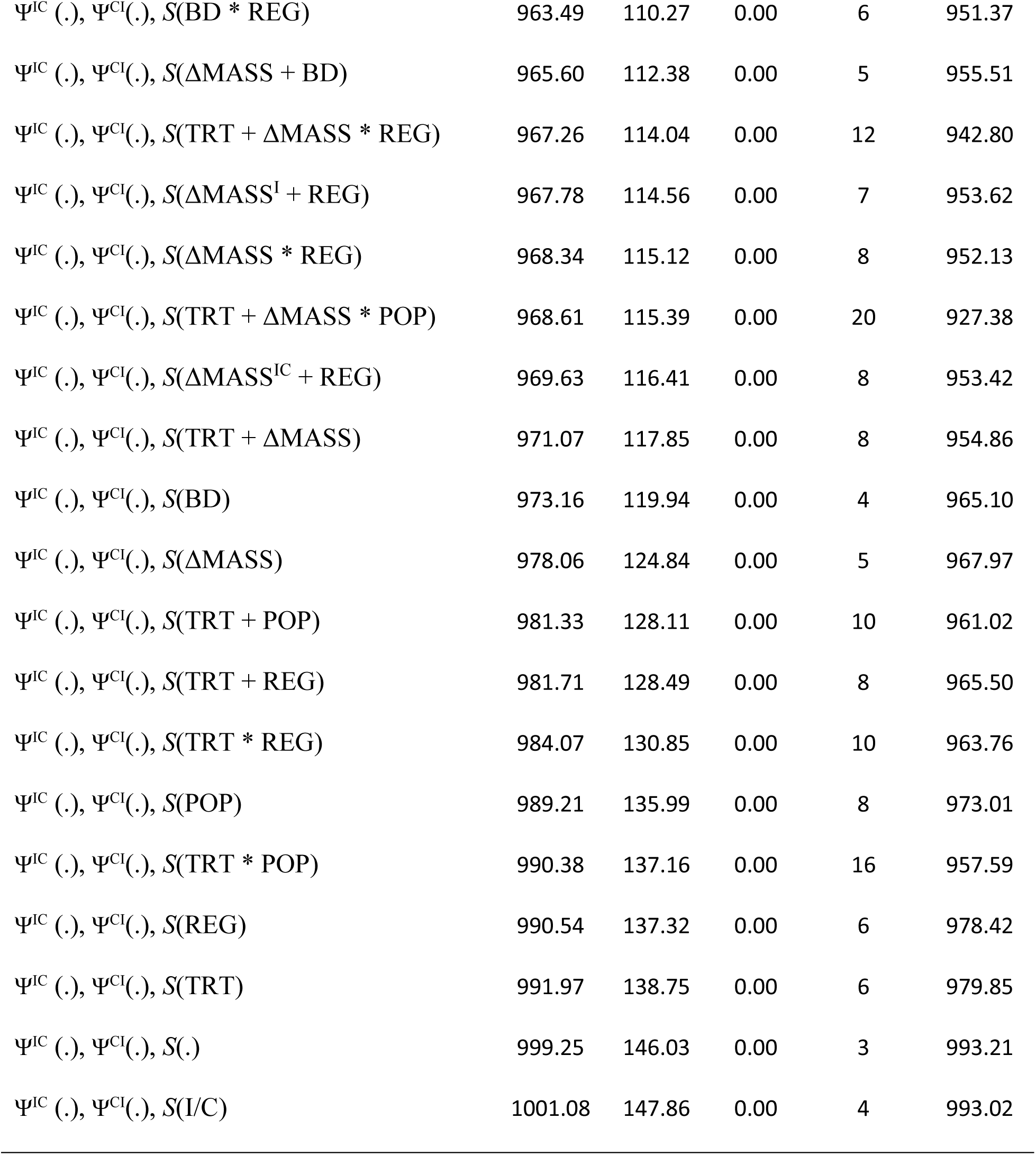

| Model | AICc | $\Delta$ AICc | $w$ | $K$ | $-2\log(L)$ |
| --- | --- | --- | --- | --- | --- |
| $\Psi^{IC}(\cdot), \Psi^{CI}(\cdot), S(BD + MASS * POP)$ | 853.22 | 0.00 | 0.89 | 14 | 824.61 |
| $\Psi^{IC}(\cdot), \Psi^{CI}(\cdot), S(BD + MASS * REG)$ | 857.45 | 4.23 | 0.11 | 8 | 841.24 |
| $\Psi^{IC}(\cdot), \Psi^{CI}(\cdot), S(BD + MASS + POP)$ | 865.23 | 12.01 | 0.00 | 8 | 849.03 |
| $\Psi^{IC}(\cdot), \Psi^{CI}(\cdot), S(BD + MASS + REG)$ | 866.71 | 13.49 | 0.00 | 6 | 854.59 |
| $\Psi^{IC}(\cdot), \Psi^{CI}(\cdot), S(MASS * POP)$ | 886.65 | 33.43 | 0.00 | 12 | 862.20 |
| $\Psi^{IC}(\cdot), \Psi^{CI}(\cdot), S(MASS * REG)$ | 890.02 | 36.80 | 0.00 | 8 | 873.82 |
| $\Psi^{IC}(\cdot), \Psi^{CI}(\cdot), S(MASS^{IC} + REG)$ | 890.06 | 36.84 | 0.00 | 8 | 873.85 |
| $\Psi^{IC} (.), \Psi^{CI}(.), S(\text{TRT} + \text{MASS} * \text{REG})$ | 891.50 | 38.28 | 0.00 | 12 | 867.05 |
| $\Psi^{IC} (.), \Psi^{CI}(.), S(\text{TRT} + \text{MASS} * \text{POP})$ | 893.76 | 40.54 | 0.00 | 20 | 852.53 |
| $\Psi^{IC} (.), \Psi^{CI}(.), S(\text{MASS}^{IC} + \text{POP})$ | 894.82 | 41.60 | 0.00 | 12 | 870.37 |
| $\Psi^{IC} (.), \Psi^{CI}(.), S(\text{MASS}^I + \text{REG})$ | 895.12 | 41.90 | 0.00 | 7 | 880.96 |
| $\Psi^{IC} (.), \Psi^{CI}(.), S(\text{MASS}^I + \text{POP})$ | 895.85 | 42.63 | 0.00 | 9 | 877.60 |
| $\Psi^{IC} (.), \Psi^{CI}(.), S(\text{TRT} + \text{MASS} + \text{REG})$ | 896.08 | 42.86 | 0.00 | 9 | 877.82 |
| $\Psi^{IC} (.), \Psi^{CI}(.), S(\text{TRT} + \text{MASS} + \text{POP})$ | 897.32 | 44.10 | 0.00 | 11 | 874.94 |
| $\Psi^{IC} (.), \Psi^{CI}(.), S(\text{MASS} + \text{BD})$ | 911.67 | 58.45 | 0.00 | 5 | 901.58 |
| $\Psi^{IC} (.), \Psi^{CI}(.), S(\text{MASS} + \Delta \text{MASS})$ | 924.01 | 70.79 | 0.00 | 6 | 911.89 |
| $\Psi^{IC} (.), \Psi^{CI}(.), S(\text{TRT} + \text{MASS})$ | 933.94 | 80.72 | 0.00 | 8 | 917.74 |
| $\Psi^{IC} (.), \Psi^{CI}(.), S(\text{MASS})$ | 936.80 | 83.58 | 0.00 | 5 | 926.71 |
| $\Psi^{IC} (.), \Psi^{CI}(.), S(\text{BD} + \Delta \text{MASS} + \text{POP})$ | 940.31 | 87.09 | 0.00 | 8 | 924.10 |
| $\Psi^{IC} (.), \Psi^{CI}(.), S(\text{BD} + \Delta \text{MASS} * \text{POP})$ | 945.87 | 92.65 | 0.00 | 14 | 917.26 |
| $\Psi^{IC} (.), \Psi^{CI}(.), S(\text{BD} + \text{POP})$ | 951.68 | 98.46 | 0.00 | 7 | 937.52 |
| $\Psi^{IC} (.), \Psi^{CI}(.), S(\text{BD} + \Delta \text{MASS} + \text{REG})$ | 954.46 | 101.24 | 0.00 | 6 | 942.34 |
| $\Psi^{IC} (.), \Psi^{CI}(.), S(\text{BD} * \text{POP})$ | 955.69 | 102.47 | 0.00 | 10 | 935.37 |
| $\Psi^{IC} (.), \Psi^{CI}(.), S(\text{BD} + \Delta \text{MASS} * \text{REG})$ | 956.33 | 103.11 | 0.00 | 8 | 940.13 |
| $\Psi^{IC} (.), \Psi^{CI}(.), S(\text{TRT} + \Delta \text{MASS} + \text{POP})$ | 956.38 | 103.16 | 0.00 | 11 | 934.00 |
| $\Psi^{IC} (.), \Psi^{CI}(.), S(\Delta \text{MASS}^I + \text{POP})$ | 959.56 | 106.34 | 0.00 | 9 | 941.30 |
| $\Psi^{IC} (.), \Psi^{CI}(.), S(\Delta \text{MASS} * \text{POP})$ | 959.96 | 106.74 | 0.00 | 12 | 935.51 |
| $\Psi^{IC} (.), \Psi^{CI}(.), S(\text{BD} + \text{REG})$ | 961.61 | 108.39 | 0.00 | 5 | 951.52 |
| $\Psi^{IC} (.), \Psi^{CI}(.), S(\text{TRT} + \Delta \text{MASS} + \text{REG})$ | 962.29 | 109.07 | 0.00 | 9 | 944.03 |
| $\Psi^{IC} (.), \Psi^{CI}(.), S(\Delta \text{MASS}^{IC} + \text{POP})$ | 963.22 | 110.00 | 0.00 | 12 | 938.77 |
| $\Psi^{IC}(.), \Psi^{CI}(.), S(BD * REG)$ | 963.49 | 110.27 | 0.00 | 6 | 951.37 |
| $\Psi^{IC}(.), \Psi^{CI}(.), S(\Delta MASS + BD)$ | 965.60 | 112.38 | 0.00 | 5 | 955.51 |
| $\Psi^{IC}(.), \Psi^{CI}(.), S(TRT + \Delta MASS * REG)$ | 967.26 | 114.04 | 0.00 | 12 | 942.80 |
| $\Psi^{IC}(.), \Psi^{CI}(.), S(\Delta MASS^I + REG)$ | 967.78 | 114.56 | 0.00 | 7 | 953.62 |
| $\Psi^{IC}(.), \Psi^{CI}(.), S(\Delta MASS * REG)$ | 968.34 | 115.12 | 0.00 | 8 | 952.13 |
| $\Psi^{IC}(.), \Psi^{CI}(.), S(TRT + \Delta MASS * POP)$ | 968.61 | 115.39 | 0.00 | 20 | 927.38 |
| $\Psi^{IC}(.), \Psi^{CI}(.), S(\Delta MASS^{IC} + REG)$ | 969.63 | 116.41 | 0.00 | 8 | 953.42 |
| $\Psi^{IC}(.), \Psi^{CI}(.), S(TRT + \Delta MASS)$ | 971.07 | 117.85 | 0.00 | 8 | 954.86 |
| $\Psi^{IC}(.), \Psi^{CI}(.), S(BD)$ | 973.16 | 119.94 | 0.00 | 4 | 965.10 |
| $\Psi^{IC}(.), \Psi^{CI}(.), S(\Delta MASS)$ | 978.06 | 124.84 | 0.00 | 5 | 967.97 |
| $\Psi^{IC}(.), \Psi^{CI}(.), S(TRT + POP)$ | 981.33 | 128.11 | 0.00 | 10 | 961.02 |
| $\Psi^{IC}(.), \Psi^{CI}(.), S(TRT + REG)$ | 981.71 | 128.49 | 0.00 | 8 | 965.50 |
| $\Psi^{IC}(.), \Psi^{CI}(.), S(TRT * REG)$ | 984.07 | 130.85 | 0.00 | 10 | 963.76 |
| $\Psi^{IC}(.), \Psi^{CI}(.), S(POP)$ | 989.21 | 135.99 | 0.00 | 8 | 973.01 |
| $\Psi^{IC}(.), \Psi^{CI}(.), S(TRT * POP)$ | 990.38 | 137.16 | 0.00 | 16 | 957.59 |
| $\Psi^{IC}(.), \Psi^{CI}(.), S(REG)$ | 990.54 | 137.32 | 0.00 | 6 | 978.42 |
| $\Psi^{IC}(.), \Psi^{CI}(.), S(TRT)$ | 991.97 | 138.75 | 0.00 | 6 | 979.85 |
| $\Psi^{IC}(.), \Psi^{CI}(.), S(.)$ | 999.25 | 146.03 | 0.00 | 3 | 993.21 |
| $\Psi^{IC}(.), \Psi^{CI}(.), S(I/C)$ | 1001.08 | 147.86 | 0.00 | 4 | 993.02 |

